# Mapping concept and relational semantic representation in the brain using large language models

**DOI:** 10.1101/2025.06.12.659304

**Authors:** Paul C. Bogdan, Roberto Cabeza, Simon W. Davis

## Abstract

How the brain organizes semantic information is one of the most challenging and expansive questions in cognitive neuroscience. To shed light on this issue, prior studies have attempted to decode how the brain represents concepts. We instead examined how relational information is encoded, which we pursued by submitting texts to a contemporary large language model and extracting relational embeddings from the model. Using behavioral data (N = 636), we found these embeddings capture independent information about scenes and objects, along with relational information on their semantic links. Turning to fMRI data (N = 60), we leveraged these embeddings for representational similarity analysis: The occipitotemporal cortex represents concepts in isolation, whereas the dorsolateral prefrontal cortex and basal ganglia principally encode relational information. Relational coding within prefrontal and striatal areas also tracks how participants reason about scenes and objects. Altogether, this research maps how information progresses from concept-level to integrative forms and how this translates into behavior.

## 1. Introduction

Humans make sense of the world by mapping sensory information to concepts, which are embedded within complex semantic networks describing said concepts and their relations.

Understanding the structure of these semantic networks has long been a central goal of cognitive neuroscience. Network structure can be modeled from human ratings of concepts’ similarity,^1,2^ from examining concepts’ overlapping features^3^, or increasingly from analysis of concepts’ positions within the semantic spaces defined by machine learning algorithms.^4–7^ For example, word embeddings for “*football*” and “*basketball*” gathered from a tool like word2vec may indicate that these two items, due to sharing common contexts and functions, are semantically similar.^7^ If a brain region likewise displays similar activation patterns when viewing a football or a basketball, then one can infer that the region encodes semantic information. This approach to studying neural information, formalized as representational similarity analysis (RSA), has proven transformative for clarifying the organization of concepts in the brain.^8^ However, most representational studies overlook a crucial aspect of semantics: the mechanisms by which the brain encodes the multifaceted semantic *relationships* between concepts – e.g., the information common across egg-chicken and seed-tree relationships. Abstract relationships like these go beyond generic similarity and differ in fundamental ways from the types of concept-level information typically investigated by studies on representation. Yet, relational information is just as (if not more) fundamental to semantic processing and executive reasoning in humans.^9,10^

A comprehensive understanding of the brain’s coding requires considering the different forms of information represented. Significant progress has been made in clarifying concept representation – e.g., RSA demonstrates that ventral stream (occipitotemporal) regions robustly encode perceptual and semantic features of visual stimuli.^11–13^ By contrast, work on relational coding is less developed, and the results on this topic are less precise. Associative memory research shows that heightened hippocampal and medial entorhinal activity when viewing a pair of stimuli strengthens binding and improves recall of which items appeared together.^14–16^ However, it remains unknown what mechanisms humans use to encode the semantics of *how* items are related. Semantic analogy tasks approach this issue, and meta-analyses on such tasks show that relational reasoning specifically upregulates dorsolateral prefrontal cortex (dLPFC) and basal ganglia (BG) (caudate head) activity.^17^ Tasks requiring reasoning about spatial and numerical relations further engage the angular gyrus and intraparietal sulcus.^18,19^ These different regions may coordinate to support relational processing via frontal-basal-thalamic loops and cortical networks,^20–24^ although none of this earlier work has attempted to model relational information as done by RSA studies mapping concept representation. This gap limits the differentiation of regions encoding relational information (multivariate effect) from those facilitating it (univariate effect).

We propose that modern large language models (LLMs) offer an ideal tool to study how people represent the relationship between concepts. Unlike word-embedding methods, which are based on words’ usages within small context windows (e.g., word2vec or GloVe), contemporary LLMs are trained on longer texts and rely on a transformer architecture that models word interactions in a generalizable manner. LLMs accomplish this using ‘attention’ layers that compute how two words are related and move information between words. When processing a text like “*In the pavement, a weed*,” an LLM encodes not only the independent meaning of *pavement* and *weed*, but also how *pavement* is relevant to *weed* – e.g., weeds exist in the cracks of pavement, which is a relational feature that could be similarly expressed for “*In the city, a rat*”. The complete *pavement*-*weed* relation can be seen as a collection of such relational features – a distributed coding scheme describing LLM processing,^25^ which parallels the coding seen in human semantic networks.^3^

LLMs will simultaneously encode a text’s concept-level and its relation-level information. This dual function is both a strength and a weakness. Because of this multifaceted processing, prior studies have found strong correlations between LLMs’ internal states and the brain data of humans reading lengthy texts.^26–28^ However, this approach will also create ambiguity, as these prior studies’ results reflect an unclear mix of concept and relational representation, severely limiting neuroscientific conclusions.

The present research aims to distinguish the coding of concepts and relations, which we accomplish by employing an original approach to LLM-based analysis. To isolate the behavioral and neural representations of relations, independent of their constituent objects, we use an LLM to parse standardized statements mentioning just two stimuli. We extract the LLM’s internal states to produce embeddings, which we distill to only relational information. For instance, to model the *pavement-weed* semantic relation, we generate an embedding for “*In the pavement, a weed*” and effectively subtract concept vectors for *pavement* and *weed* alone. This strategy is inspired by earlier work subtracting features from embeddings.^29,30^ Below, we show how this strategy eliminates all measurable concept-level information from the embedding while retaining its relational content. RSA can then be done with either pure concept or pure relational embeddings to disentangle these semantic dimensions.

Our investigation proceeds in two phases. In *Study 1*, we examined conceptual representation and first analyzed behavioral data to validate that embeddings generated using a modern LLM accurately capture human-reported concept semantics. We next conducted RSA using the LLM concept embeddings and fMRI data from participants viewing object stimuli in isolation, expecting to replicate extant work on the ventral stream’s role in concept representation. In *Study 2*, we focused on relational representation. We begin with behavioral analysis to validate that embeddings based on LLMs capture human-reported relational information. We expect that modern LLMs will impart particularly large improvements in modeling relational information compared to older techniques like word2vec or BERT, pointing to LLMs fundamentally changing modeling capabilities rather than being incremental improvements. We then performed RSA using LLM-based embeddings for scene-object relations, applied to fMRI data from participants who reasoned about pictures of scenes and objects. We expected to identify brain regions that principally encode relational content and differ from those leading concept coding. We expected to additionally identify areas where the strength of relational representation predicts participants’ judgment in the scene-object task (e.g., stronger representation predicts improved task performance), linking relational coding to behavior. By examining these different aspects of information processing, we aim to shed light on how the brain’s different areas cooperate to support general semantic processing.

## 2. Results

### 2.1. Study 1. Mapping concept representation in the brain

Our first study investigated concept representation and is divided into two parts: Study 1A evaluates the efficacy of our LLM approach for defining concept embeddings. We specifically measured the correspondence between embeddings generated via Llama 3.2-3B and an existing database of propositional concept features developed through human studies. Study 1B then examined fMRI data from tasks where participants viewed objects in isolation. The results are expected to outline the structure of concept representation.

#### 2.1.1. Study 1A. Behavioral analysis and validation of LLM-based concept embeddings

We first evaluated concept-level semantics using public data from a normative study on object features (https://mariamh.shinyapps.io/dinolabobjects/).^31^ In the study, 566 participants saw pictures of 995 everyday objects and were asked to generate properties (or features) describing each image (e.g., shown an *apple*, a participant may respond “*is edible*”). For the present analysis, a simple text was crafted for each object containing only an article/determiner and the object’s name (e.g., “*An apple*”). Each text was submitted to the open-source LLM, Llama 3.2-3B (28 layers). Embeddings were defined as the LLM’s residual stream activity at the final token position, computed either for single layers or while pooling multiple middle layers (layers 4-16). Using the embeddings, support vector machines (SVMs) were separately fit for each of the fifty most common features in the dataset, predicting whether a given object had (1) or did not have (0) said feature (**Figure 1A**). The resulting accuracies were compared to those of SVMs fit based on objects’ word2vec embeddings. For fair comparisons, the LLM embeddings were reduced to 300 dimensions (same as word2vec) using principal component analysis before classification.

**Figure 1.**
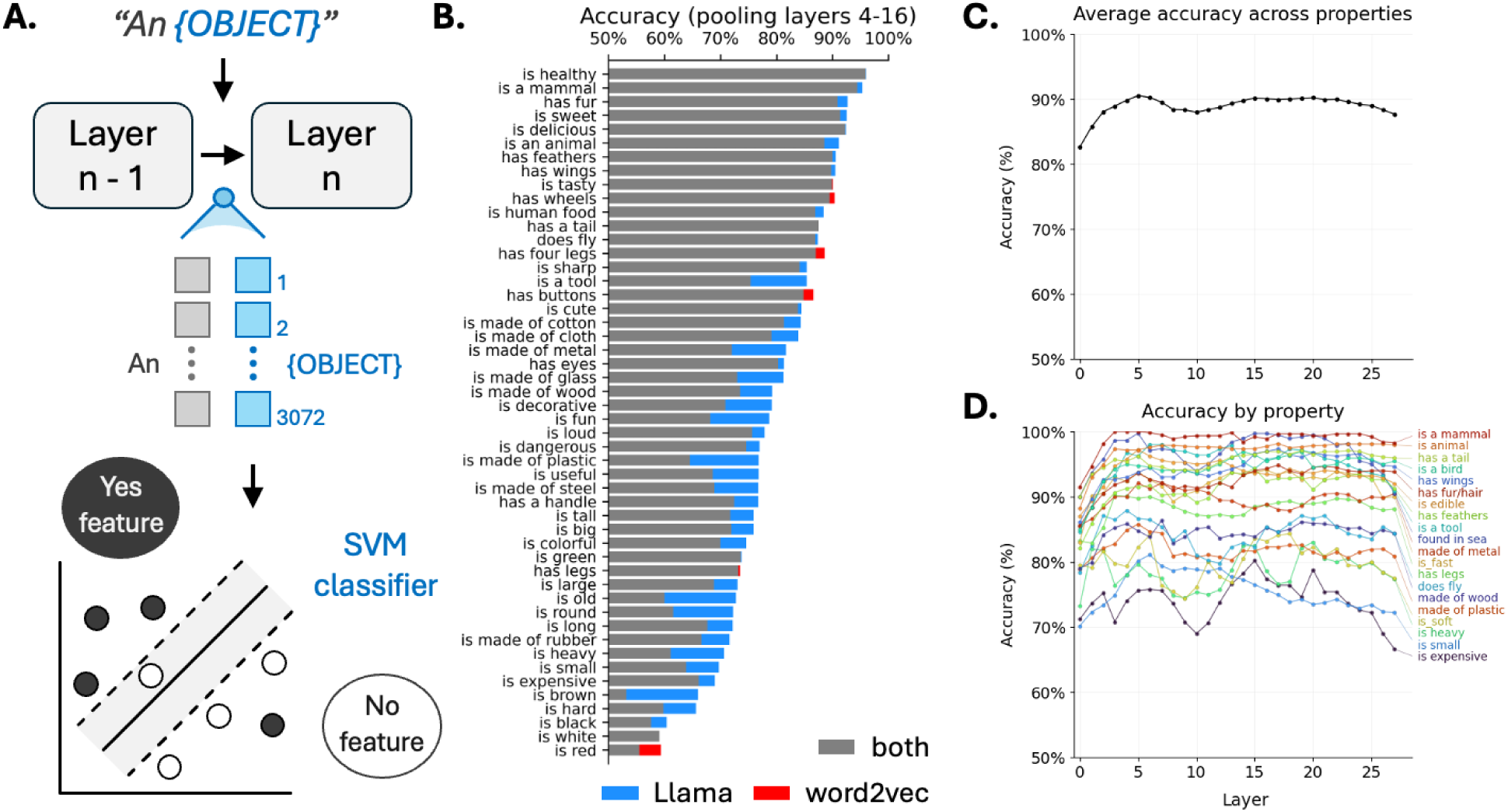
Representation of isolated item semantics by LLMs. **A.** Illustration of the approach, where short item texts were submitted to Llama 3.2-3B, and embeddings were extracted from the residual stream at the last token position. For the fifty most common features in the dataset, separate SVMs were fit to predict whether a particular object has or does not have a given feature. Accuracy was evaluated using 10-repeated 5-fold cross-validation while omitting objects randomly so that chance accuracy is 50%. As a comparison, classification was also tested using word2vec embeddings. **B.** The bars here represent the cross-validated accuracy for each of the fifty features, using embeddings that concatenate layer 4-16 activity. Blue indicates an accuracy advantage by Llama 3.2-3B over word2vec, whereas red indicates the opposite. **C.** Accuracy is shown, averaged across the different features using embeddings based on singular layers. **D.** Layer-by-layer accuracy separately for each feature is presented here; only twenty features are shown for visual clarity.

The LLM embeddings predicted nearly every feature at above-chance accuracy (mean [M] = 79.5%), consistently surpassing the accuracy of word2vec-based predictions (M = 75.6%) (**Figures 1B-D**). The Llama 3.2-3B accuracy also consistently surpassed predictions based on embeddings from a historic transformer model (BERT), tested with the same procedures (M = 69.0%). Contemporary LLM embeddings thus robustly capture concept-level semantics.

This correspondence with human-reported semantic information further suggests that LLM embeddings may be more interpretable and cognitively meaningful than embeddings based on older techniques.

We conducted similar tests using alternative LLMs, which surpassed the accuracies of word2vec and BERT: Llama 2-7B (M = 79.4%), Llama 3.3-70B-Instruct (M = 79.6%), Qwen 2.5-72B (M = 78.6%), and DeepSeek V2-Lite (16B; M = 78.8%). These alternative models are all based on transformers but differ in size and other implementation or training details. The consistently high accuracy levels show how accurate semantic representation is a general property of modern LLMs. We proceed with Llama 3.2-3B, as it is the leanest, can be readily run on a consumer computer, and appears sufficient for high-quality concept modeling, positioning it well for cognitive and neuroscientific research.

#### 2.1.2. Study 1B. Neural analysis of concept representation using objects

Having demonstrated the strength of LLM-based embeddings for modeling human-reported semantic features, we turn to analyzing the representation of concepts in the brain. Here, we rely on fMRI data collected during each task of a multi-stage experiment on how humans encode conceptual and relational information. The stages consisted of: (1) an object naming task, (2) a task where participants judged the link between an object and a scene, (3) a conceptual recognition task for object memory, and (4) a visual recognition task for object memory. Study 2 will later focus on relational representation in the second task, which is the most unique element of the design and makes the dataset well-suited for our overarching aims of mapping concept and relational representation. However, for Study 1B, we focused on just tasks 1, 3, and 4, where participants viewed isolated objects, and we use the fMRI data generated from these tasks to examine single concept representation.

Standard RSA was first performed with the embeddings generated by submitting isolated object texts (“*An {object}*”) to Llama 3.2-3B (**Figure 2A**). Based on these embeddings, across-trial correlations were used to construct a 114 × 114 representational similarity matrix (RSM) for each participant and task stage. Corresponding neural similarity matrices (NSMs) were produced for 246 Brainnetome-atlas regions of interest (ROIs) by correlating voxel-wise activity across trials.

**Figure 2.**
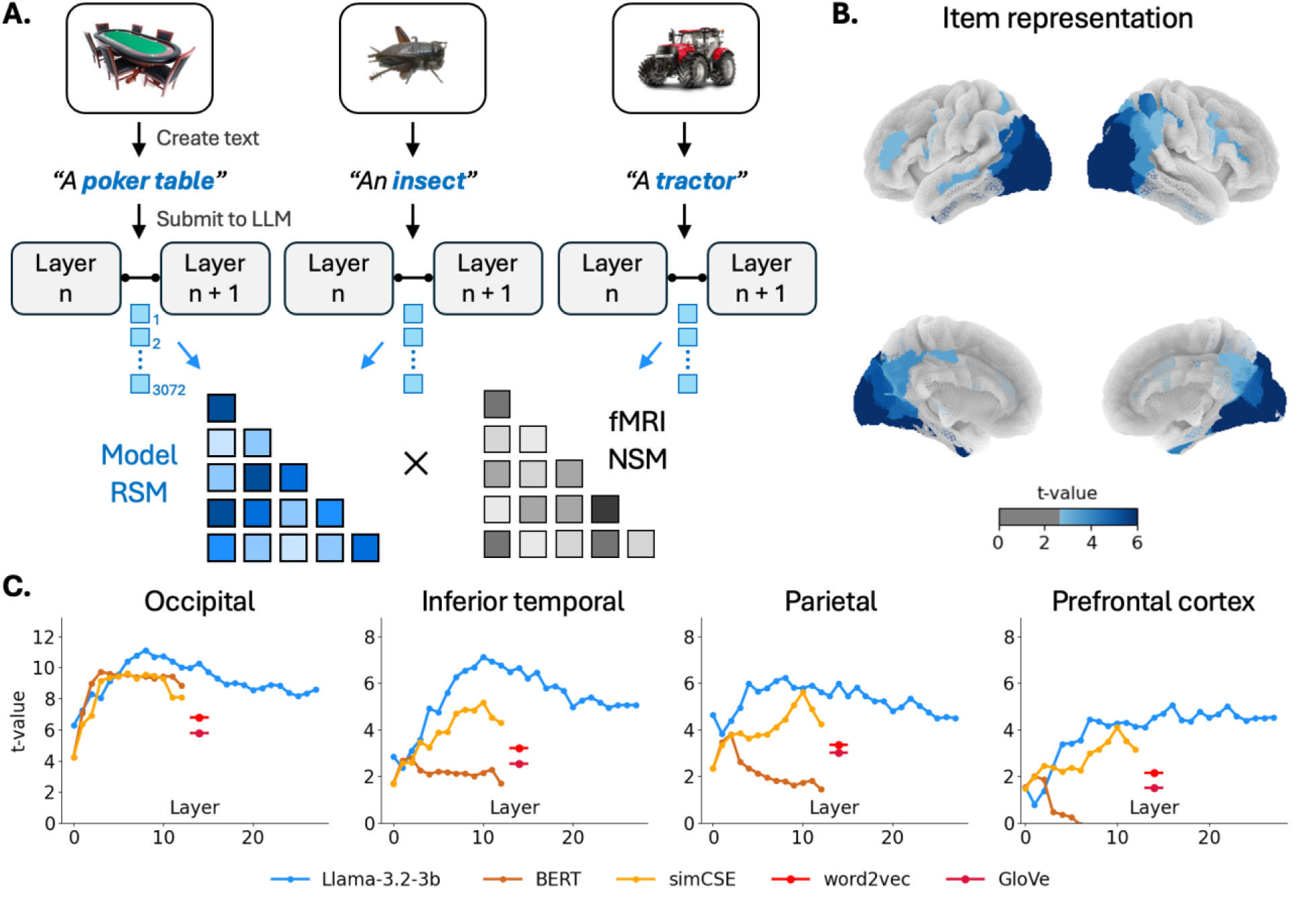
Single concept semantics RSA. **A.** For each of the 114 objects presented in the task, a text was prepared and submitted to Llama 3.2-3B. The LLM’s residual stream was extracted and concatenated across layers 4-16 to produce embeddings. Across-trial correlations between the embeddings were computed to form model RSMs, and corresponding fMRI NSMs were formed via correlations between trials’ voxel-wise activity patterns. A separate NSM was produced for each Brainnetome atlas ROI (246 ROIs in total). Second-order RSM x NSM correlations were computed, averaged across the three task stages by participant, then submitted to a group-level one-sample t-test. **B.** This t-test yielded significant effects in the ROIs here; all results are false-discovery rate corrected for 246 tests (p_corrected_ <.05). **C.** Layer-by-layer RSA results are also provided for four large-scale areas (NSMs computed based on pooling voxels among all constituent ROIs; see Methods for anatomical definitions).

For each ROI, second-order RSM × NSM Spearman correlations were next calculated for all sixty participants and three isolated-object task stages (180 correlations). These correlations were next averaged by participant (60 mean correlations). For group-level analysis of each ROI, those sixty points were submitted to a one-sample t-test. This yielded strong RSA effects throughout the occipital and inferior temporal lobes – areas known to represent objects (**Figure 2B**).^11–13^ The shown results are based on LLM embeddings that pool across layers 4-16, although examining single-layer embeddings likewise showed that effects were nominally strongest in the occipital and inferior temporal lobes (**Figure 2B**). These LLM-based RSA effects were notably stronger than the RSA results associated with older models, such as word2vec, GloVe (a variant of word2vec), BERT, or SimCSE (a variant of BERT) (**Figure 2C**).

Note that we have pooled data from all three isolated object tasks to maximize statistical power and because we do not have any hypotheses specific to any specific task. However, analyses of solely, for instance, the object-naming task yielded similar results: significant (*p_corrected_ <.05*) effects across 33 occipitotemporal ROIs and 17 parietal ROIs, while effects elsewhere are sparse. Overall, the strong LLM patterns and consistency with the established RSA literature on object processing^11–13^ speak to the validity of the LLM approach and set the foundation for more diverse and novel directions.

### 2.2. Study 2. Mapping relational representation in the brain

Study 2 focused on the representation of relational semantics, and like above, this study is divided into two parts: Study 2A evaluated the capacity for LLM-based embeddings to capture this relational information using the behavioral data from the scene-object task from the multi-stage experiment mentioned previously. Study 2B then applied these embeddings for mapping the brain regions responsible for encoding relational information. The brain investigation focuses on contrasting the regions performing relational coding from those noted above on concept coding.

#### 2.2.1. Study 2A. Behavioral analysis and validation of LLM-based relational embeddings

Our behavioral analyses focus on a task where participants reasoned about a scene and object. Specifically, participants were shown a picture of a recognizable scene followed by an object (e.g., a *farm* followed by a *tractor* or a *bathroom* followed by a *chainsaw*). For each pair, participants rated the likelihood of finding the object in the scene using a 4-point scale (1 = “*Very unlikely*”, 4 = “*very likely*”). The task was designed to elicit the full ratings by including scene-object pairs selected to be incongruent, neutral, or congruent (one-third each; see Methods). In total, 60 participants each rated 114 scene-object combinations (342 unique pairs overall).

The present analyses used LLM embeddings to predict each scene-object pair’s mean likelihood rating, averaged across subjects. Scene-object embeddings were created by submitting two crafted texts (“*At the {scene}, an {object}*” and “*An {object} was in a {scene}”*) to Llama 3.2-3B (**Figure 3A**); see Methods for discussion and additional results of text crafting. The LLM’s residual stream was extracted at the final token positions, concatenated across layers 4-16, and averaged across the two texts. The resulting embeddings were submitted to a ridge regression predicting the average human-reported pair likelihood; a ridge regression was used, rather than a standard linear regression, as there are more embedding dimensions (36,864) than examples (342), which would saturate a standard regression.

**Figure 3.**
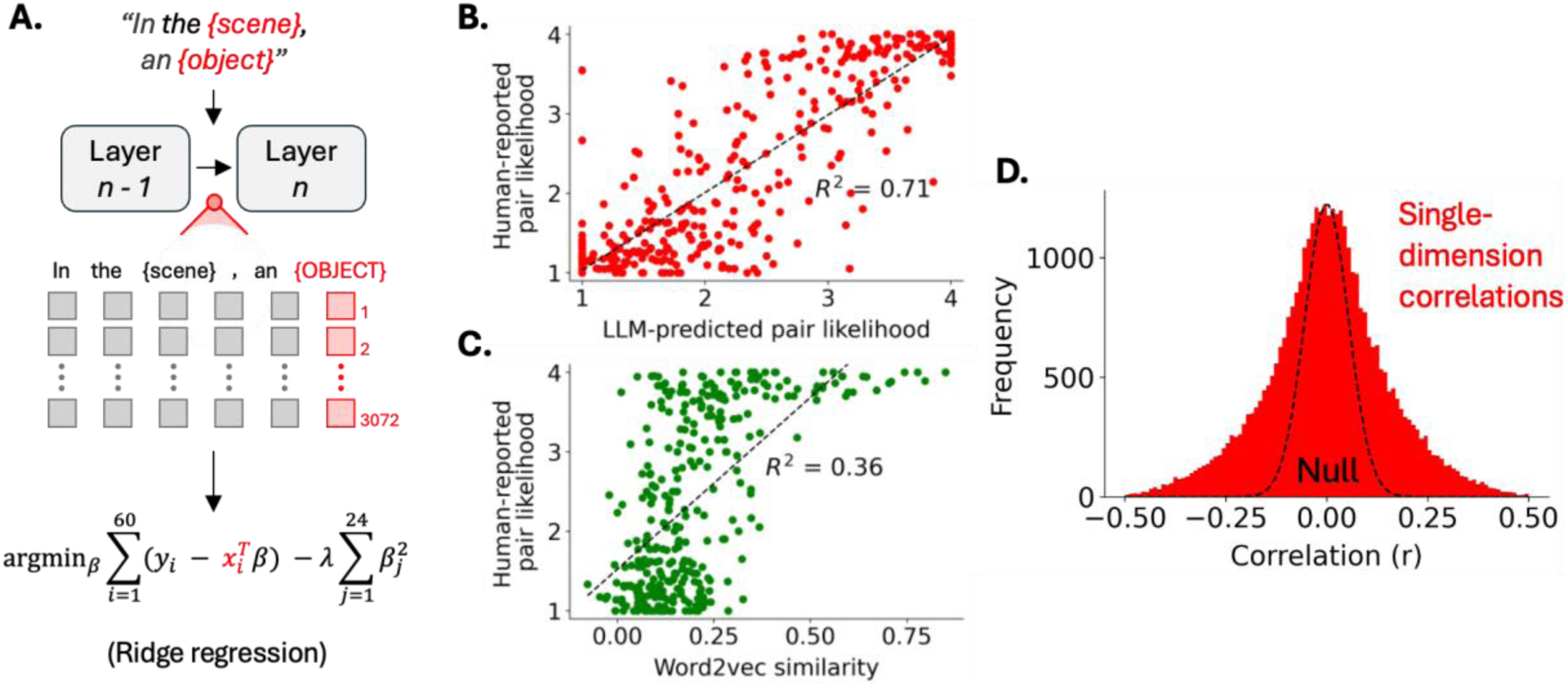
Representation of relational semantics by LLMs. **A.** Illustration of the approach, where simple scene-object texts were submitted to Llama 3.2-3B, and embeddings were extracted from the residual stream at the last token’s position, concatenated across layers 4-16. **B.** The embeddings strongly predicted scene-object pairs’ associated human-reported likelihood of finding the object in the scene. **C.** Similarity between the scene and object word2vec embeddings less predicted similarity. **D.** The correlations between human-reported likelihood and each of the LLM’s 36,864 states’ activations (12 layers × 3072 internal states per layer) are shown here as a histogram. For reference, the distribution of correlations that would arise from random data is shown by the dashed line (a zero-centered normal distribution).

The LLM embeddings strongly predicted scene-object ratings (*R*^2^ =.71; **Figure 3B**). This rate far surpassed what older transformer models like BERT achieve (*R*^2^ =.07) and also exceeded predictions based on the correlational similarity between *scene* and *object* word2vec embeddings (*R*^2^ =.36; **Figure 3C**). Note, word2vec’s disadvantage does not stem from non-normality in the data, as a Spearman correlation between word2vec similarity and scene-object ratings yielded a weaker effect (*ρ* =.58, *ρ*^2^ =.33). Word2vec’s disadvantage also does not stem from it being a scalar, and submitting the 300-element difference between object and scene vectors to a ridge regression produced null predictions (*R*^2^ < 0). Thus, we have strong evidence that the embeddings from a modern LLM uniquely and robustly describe the relatedness between scenes and objects.

The LLM embeddings’ accurate predictions are rooted in multivariate relational information. That is, scene-object likelihood positively correlated with activity in some LLM dimensions but negatively correlated with others (**Figure 3D**). Statistically, this occurred significantly more often than expected from a zero-centered null distribution of correlations (Kurtosis: K = 0.64, *p* <<.001). This pattern can be explained by different directions in the embedding describing different ways that two items can be related. Preliminary exploratory analyses showed this using a toy dataset producing embeddings of crafted texts “*An {animal} and a {food}*”. An SVM trained on the embeddings can significantly predict the *a priori* relational feature “*would eat*” (1 for a carnivore with meat or an herbivore with food and 0 otherwise).

Hence, this narrow relational proposition is expressed by the LLM. A full embedding may describe many relational features like these, much like how the dimensions of an item embedding describe different item features.

#### 2.2.2. Study 2B. Neural analysis of relational representation

RSA was conducted using the fMRI data collected during the aforementioned scene-object task. In each trial, participants viewed a scene and then an object, and RSA was conducted during the screen where the object was shown (**Figure 4A**). Before conducting tests using relational embeddings, we performed concept-level RSA for object and scene representations. Concept-level RSA for just the object’s representation (*“An {object}”*) yielded ventral stream results like those identified in Study 1B (**Figure 4B**). Concept-level RSA was also done for the presented scene’s representation (“*A {scene}”* embedding); note that the brain measurements are based on the screen where the object was shown, and the present analysis thus taps into how participants maintained the scene representation in working memory. Focusing on scene representation implicated new regions: namely, the inferior parietal lobule (IPL) along with medial parietal structures, such as the retrosplenial cortex, now showed the strong effects (**Figure 4B**). Additionally, the dLPFC and middle temporal gyrus (MTG) emerged, pointing to the involvement of the frontoparietal control network (FPCN). This network is often characterized by this triplet of regions (IPL-LPFC-MTG) involved in working memory.^32–34^ Yet, it remains unanswered how people specifically represent the scene-object relation.

**Figure 4.**
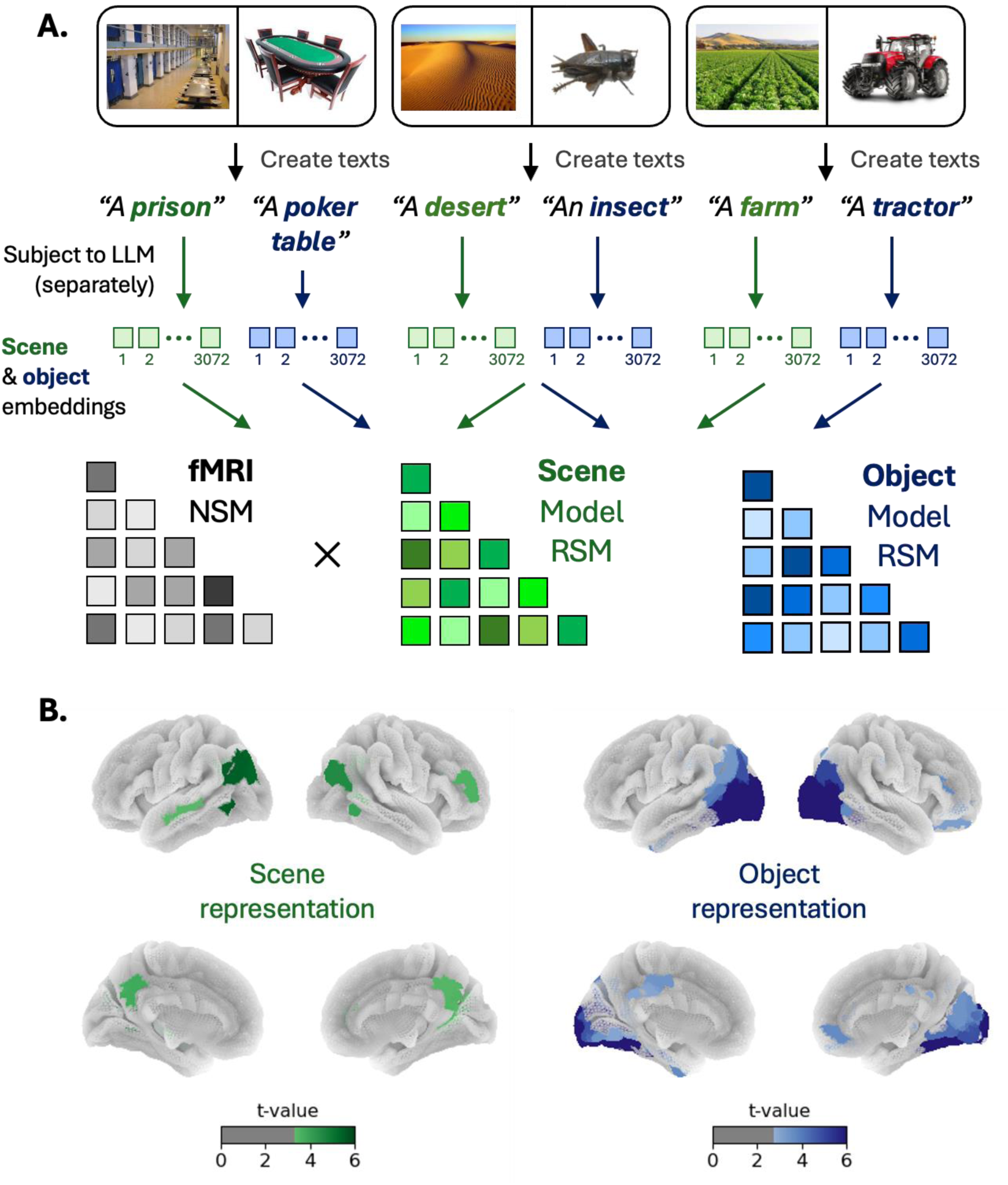
Scene and object item semantics RSA. **A.** Concept-level RSA was performed as in Study 1B, but now for scene and object representations of the scene-object task. **B.** Scene RSA and object RSA were performed independently, generating the results shown here (one-sample t-tests evaluating whether the average second-order correlation across participants is positive); shown results are false-discovery rate corrected (p_corrected_ <.05). Note that the task was designed such that a third of scene-object pairs were incongruent (left pair, neutral, or congruent. This aspect of the task is further investigated later in the results (see also Methods).

To understand how the brain parses scene-object relationships, we submitted to Llama 3.2-3B crafted texts involving both the scene and object, as in behavioral Study 2A: “*At the {scene}, an {object}*” and “*An {object} is at the {scene}*” with LLM responses averaged across the two texts to form an embedding (**Figure 5A**). To eliminate concept-level information, the embedding was mean-centered relative to the two other scene-object embeddings involving the same object – e.g., the mean of the *desert-table* and *farm-table* embeddings was subtracted from the *prison-table.* The same was also done with respect to the two other object-subtracted embeddings with the same scene. To confirm that the relational embeddings contain no concept-level information, we returned to the fMRI data from Study 1B on the isolated object tasks. These tasks presented no scenes, so RSA using these relational embeddings should produce no significant effects. This is indeed the case: all 246 ROIs yielded *p_uncorrected_ ≥*.05, suggesting that there may even be a slight bias away from single-concept effects. Hence, our mean-centering strategy removes all measurable concept-level information and produces a pure relational embedding.

**Figure 5.**
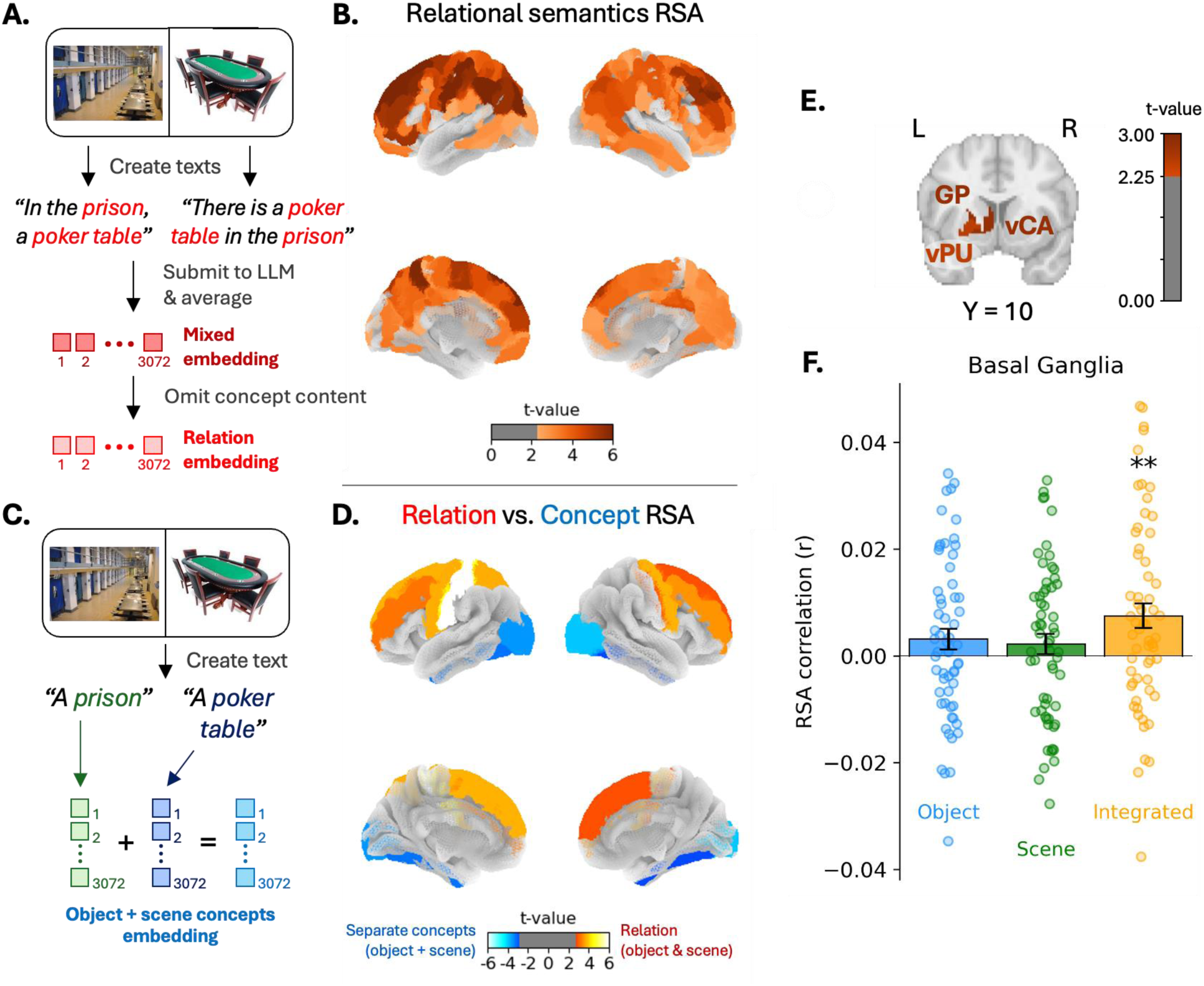
Relational semantics RSA. **A.** For each scene-object trial, two texts mentioning the scene and object were crafted and submitted to Llama 3.2-3B to produce an embedding, as for the Figure 3 relational behavior analysis. Each embedding pools data across layers 4-16 and was mean-centered with respect to other embeddings containing the same object or scene to eliminate concept-level information. **B.** The mean-centered embeddings were used for RSA, and the brain plots here show regions displaying significant relational representation per one-sample t-tests. **C.** As a point of comparison, the overall concept-level representation of the scene and object was modeled by summing scene and object concept embeddings and used for RSA. **D.** Paired t-test results are illustrated, comparing RSA effects to distinguish ROIs primarily representing relational semantics from those primarily representing single concept semantics. As this analysis is a subtler evaluation than the one-sample t-tests, ROIs’ voxels were pooled based on their Brainnetome atlas regional labels (e.g., the six L superior frontal gyrus ROIs were combined into one large ROI), which increased statistical power. Results are false-discovery rate corrected (p_corrected_ <.05). **E.** Not visualized in the prior brain plots, three basal ganglia ROIs yielded significant effects: the L globus pallidus (GP), L ventral putamen (vPU), L ventral caudate (vCA). **F.** Bar graphs show the relational-embedding RSA effect in the basal ganglia, based on pooling voxels across all twelve basal ganglia ROIs. Each dot represents the second-order correlation of one participant. **, p <.01.

Analyzing the scene-object task using relational embeddings, RSA effects were strongest in dorsal and lateral areas (**Figure 5B**), particularly in the IPL and LPFC. Notably, significant MTG representation emerged, while its neighboring superior and inferior temporal gyri displayed null effects. Together with the IPL and LPFC, this set of regions is further striking evidence for the relevancy of the FPCN. To distinguish areas principally involved in relational versus single-concept processing, the relational RSA effects were contrasted to the earlier scene and object concept RSA effects. The overall degree of concept-level representation was modeled by performing RSA with respect to the sum of isolated object (“*An {object}*”) and scene (“*A {scene}*”) embeddings (**Figure 5C**). Then, paired t-tests compared the second-order relational RSA and concept RSA correlations, which yielded a dissociation: the ventral stream specializes in parsing concept semantics while the frontal lobe predominantly parses relational semantics (**Figure 5D**).

Alongside these cortical patterns, several BG structures also showed significant relational representation (**Figures 5E & 5F**). These BG effects are peculiar, as these regions are often linked to the establishment of motor associations,^35,36^ but not typically to the encoding of semantic relational representations. The BG would be further analyzed as we shifted to examining how relational representation impacts cognition and behavior.

#### 2.2.3. Linking neural relational information to scene-object judgments

If the patterns put forth reflect relational processing as generally construed, then stronger modeled representations should be associated with greater understanding by participants and improved performance in the task. This is a stricter hypothesis than those tested thus far and examines how our modeling captures cognitive processing. To test this, we examined participants’ scene-object likelihood ratings in each trial. As mentioned in Study 2A, participants used a 4-point scale to rate how likely the presented object would be found in the presented scene. The scene-object pairs were designed to elicit high/congruent (farm-tractor), medium/neutral (prison-fork), or low/incongruent (cafeteria-rose) likelihood ratings. Hence, our present analyses quantify performance in each trial as normative fit, which was computed as the absolute difference between participants’ ratings and the expected rating for each category (high: 4; medium: 2.5; low: 1).

Predicting trial-wise outcomes requires also modeling the degree of relational representation on a trial-by-trial basis. To accomplish this, a second-order correlation was computed for each trial between its corresponding NSM row and relational RSM row – illustrated in **Figure 6A** (see also earlier work using this approach^12,37^). For each participant, a within-subject correlation was taken between their normative fit and representational strength measures in each trial. These correlations were submitted to a group-level one-sample t-test, which demonstrated several significant effects (**Figure 6B**): The regions displaying the strongest correlations between relational representation and normative fit were the left and right premotor cortex, followed by dorsal PFC, with weaker significant effects in inferior parietal cortex and the middle temporal gyrus. Complementing the earlier L BG results, relational representation in the L BG also significantly predicted normative fit (pooling across L BG ROIs: *t*[59] = 2.96, *p*_corrected_ <.05); uncorrected post hoc tests showed the effect is strongest in the L ventromedial putamen (*t*[59] = 2.58). These patterns specifically stem from relational representation, and no region yields a significant effect if the correlations are instead tested using a trial-wise measure of concept representation (all *p_corrected_* >.05). Hence, relational representation in these areas specifically contributes to a more general semantic reasoning about the dynamic relationship between items.

**Figure 6.**
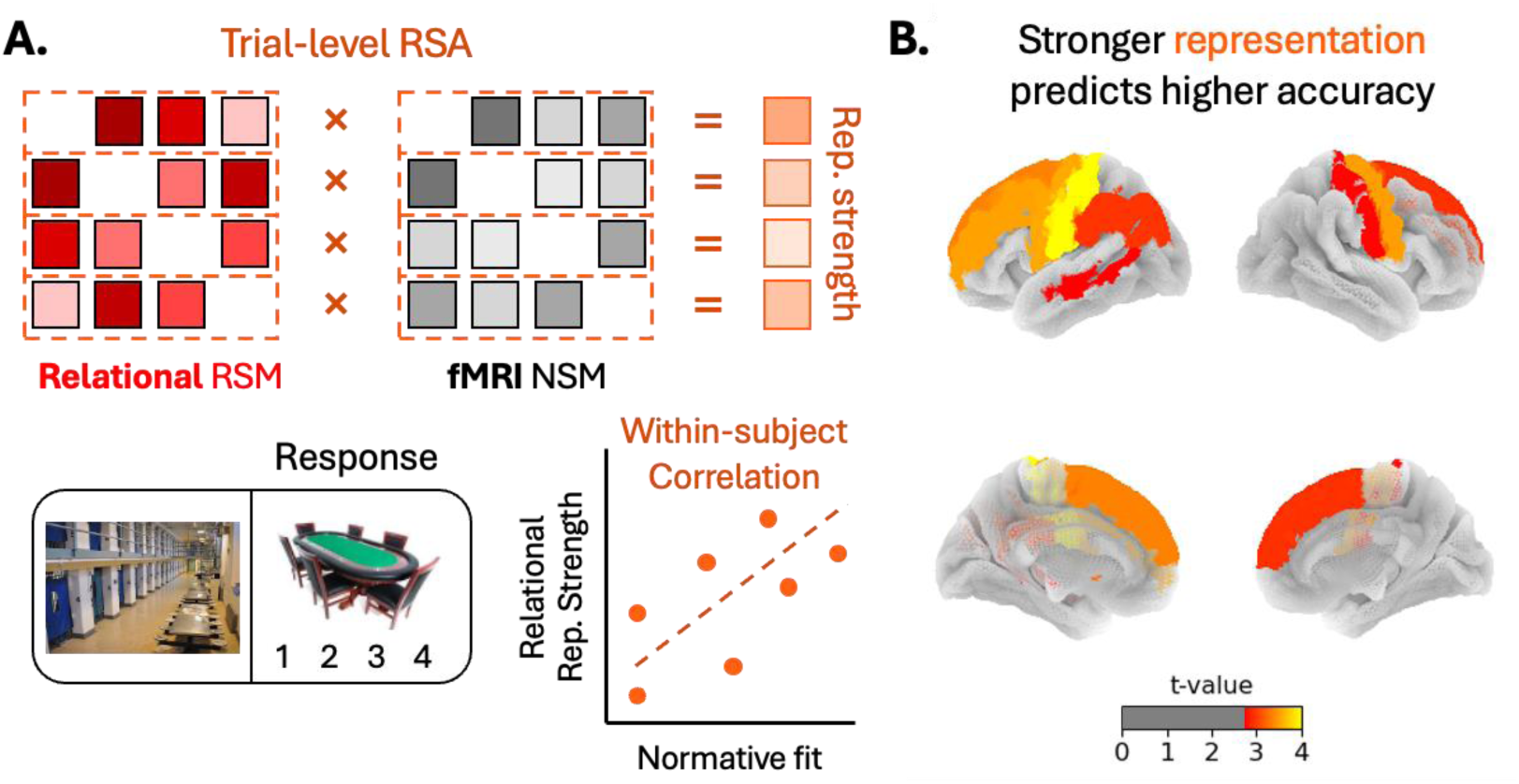
Prediction of task performance (normative fit) from relational representation. A. Trial-wise measures of representational strength were computed via second-order correlations between relational RSM and NSM rows. These were correlated within-subject with the normative fit of a participant’s response in each trial. **B.** The within-subject correlations were submitted to a one-sample t-test, identifying the ROIs where the group-average correlation was positive. Similarly to Figure 5D, the results here are presumed to be more subtle than typical RSA effects, so ROIs from the same region were pooled to form larger ROIs and increase statistical power. Shown results are false-discovery rate corrected (p_corrected_ <.05).

## 3. Discussion

The present research leverages a contemporary LLM to map how semantic information coding progresses from concept to relational representations and how this translates into participants’ relational judgments. Study 1 targets concept representation, first validating the efficacy of using LLMs’ internal states for modeling distinct aspects of conceptual information. Applying the LLM-based concept embeddings, RSA then shows how the ventral stream represents incoming objects as concepts. Study 2 next targets relational representation, first validating that the LLM embeddings robustly capture relational information. These embeddings were then leveraged for RSA. Frontoparietal regions emerged as representing relational information, particularly the frontal lobe, which represents the semantic relations between concepts more than the isolated concepts themselves. Interestingly, BG structures also contributed to parsing semantic relations. Both frontal and BG operations additionally predicted participants’ responses on the relationship between the scene and object stimuli. Altogether, these results outline how the human brain reasons about multiple objects and provide evidence supporting an analytic approach that can be generalized to other studies and questions.

Our behavioral analyses establish three methodological conclusions on the use of LLMs for modeling brain representation. *First*, semantic embeddings based on modern LLMs outperform those from simpler historic models like word2vec and BERT (**Figures 1 & 3**). High-quality modeling, in turn, enhances statistical power in all downstream analyses – evidenced further in the LLM-embeddings generating the stronger concept RSA results in the fMRI experiment than older models (**Figure 2**). *Second*, because the behavioral analyses all focused on the degree of correspondence with human reports of semantic information, the results indicate that modern LLMs effectively capture how humans themselves specifically process semantics. LLM embeddings can therefore be taken to be more interpretable than older methods, at least for the purposes of psychological and neuroscientific research. *Third*, whereas prior fMRI studies employing LLMs have focused on verbal experiments where participants read or listened to lengthy texts,^26,28^ we demonstrate how texts can be crafted to allow high-quality semantic modeling in virtually any study, even ones using pictures. Text crafting notably introduces researcher degrees of freedom, but our preliminary analyses show that minor changes to the texts minimally influence the results, so long as proper grammar is maintained (see Methods Section 4.2.2). Altogether, these three points make a strong case for LLMs to be a valuable technique for the toolbox of any researcher interested in semantic representation.

Using LLMs to model semantic information, our work puts forth several fMRI findings. Applying concept-level embeddings (“*An {object}*”), we show how the ventral stream – i.e., the early visual cortex, lateral occipital, fusiform gyrus, inferior temporal gyrus, and parahippocampal gyrus – specializes in parsing actively presented objects (**Figures 2B & 4B**). Although the relevance of the ventral stream to object processing is well established,^11–13^ our work additionally shows how the ventral stream minimally engages in any form of semantic relational representation (**Figure 5D**). This is interesting, as memory research suggests that anterior portions of the ventral stream help encode associations and semantic networks.^14,38,39^ Perhaps reconciling these ideas is that our associative modeling focused on established relations available to an LLM – e.g., knowledge about the link between a tractor and a farm. In contrast, associative memory paradigms typically employ novel associative learning and may therefore involve mechanisms orthogonal to accessing established relations. More generally, anterior ventral regions may encode *that* two concepts are related but not *how* they are related.

The IPL and medial parietal areas (retrosplenial cortex and precuneus) may operate as a bridge between concept and relational information coding. These dorsal regions robustly represent concept-level information about objects and scenes (**Figures 2 & 4**) while also engaging in relational processing (**Figure 5B**). The relational results are based on RSA using embeddings that describe information on the semantic interaction between the scene and object (e.g., “*At the {scene}, an {object}*”). Neither the IPL nor any medial parietal structure showed any clear preference for concept or relational representation (see null differences in **Figure 5D**). A dual coding role for the IPL is consistent with classical models of language, which suggest that the multi-modally connected IPL (or angular gyrus) is a key hub for thematic or combinatorial semantics.^40^ This role positions the IPL ideally to contribute to relational processing, which by its nature implies combining information from potentially disparate concepts. Likewise, the retrosplenial cortex is thought to play an integrative role in encoding spatial information.^41,42^ As bridges, these parietal structures may convey conceptual information to areas that encode solely relations, such as the LPFC.

Our RSA results on relational coding consistently show effects in the LPFC, IPL, and MTG (**Figures 5B & 6B**). This set of areas is consistent with the canonical frontoparietal control network.^32–34^ However, there is heterogeneity within this set. Whereas the IPL represents both concepts and relations, the frontal lobes process information principally at a relational level and minimally represent concepts. This is consistent with the abundance of work illustrating the PFC’s role in executive tasks,^40^ and research identifying representation in the PFC only when dealing with concepts’ abstract properties (e.g., the PFC represents items’ category memberships or their affordances^43,44^). Our work generalizes this idea across semantic relational processing broadly and shows what computations occur elsewhere in the brain to possibly support these PFC operations.

We additionally found that relational informational operations in the LPFC bear on participants’ behavior in the task. Specifically, stronger representations of the scene-object relation, particularly in areas of the PFC, predict more normative decisions about the scene and object (**Figure 6**); normativity effectively serves as a measure of task accuracy. The PFC is generally associated with semantic control functions in neuropsychological research,^45^ functional neuroimaging studies of healthy volunteers,^46^ and neuromodulation research.^47^ The current design did not explicitly manipulate executive control or semantic decision difficulty, so it is difficult to unpack how relational processing interacts with the mechanisms monitoring this processing.

Nonetheless, our more basic conclusion remains robust: participants’ judgments are linked to relational rather than concept-level informational codes in the PFC.

Notably, alongside frontoparietal effects, relational-coding RSA revealed that multiple BG structures represent relational information (**Figures 5E & 5F**) and relational representation in the BG predicts normative task responses. The consistent emergence of the basal ganglia across these two separate relational analyses may be surprising. Research on object representation generally focuses instead on cortical areas, a tendency supported by the neuropsychological literature.^48,49^ Potentially, the incorporation of BG structures in relational processing emerges from frontal-basal-thalamic loops.^20–22^ The PFC projects information to the BG, putting forth different candidate thoughts and actions, which the BG selects among through competition mechanisms.^50,51^ In turn, projections back to the PFC update working memory via gating mechanisms – e.g., a participant may hold a representation of a *poker table* concept and a *prison* concept in their PFC, then reason about how gambling can be dangerous in a prison environment via frontal-basal-thalamic loops modifying active working memory states.^52^ Such mechanisms may produce the decodable fMRI patterns generating the relational representation effects seen here.

In sum, shifting from univariate to multivariate informational analyses transformed how cognitive neuroscience studies concepts. By leveraging LLMs, the present research presents an analogous information-level approach to modeling relational coding. Together with the high-quality semantic modeling afforded by LLMs broadly increasing statistical power, it becomes possible to map the progression of information processing between single-item and multi-item relational stages. We apply this strategy for fMRI analysis and show how most regions of the brain represent information in some way, and how this coalesces to influence human behavior and inferences about items’ links. This focus uncovered effects in areas not traditionally thought to encode stimulus information – namely, basal ganglia structures – but whose roles in information processing emerge when examining information at the relational level. We hope that the findings and general methodological principles put forth will encourage future work pursuing a more multifaceted understanding of information processing.

## 4. Methods

### 4.1. Study 1A concept data and analysis

#### 4.1.1. Concept data

The investigation of concept-level semantics relied on data previously collected by our group, where 566 participants were each shown pictures of 40 items/concepts (e.g., birds, buildings, tools). The items varied between participants, such that 995 concepts were shown in total. For each concept, each participant was asked to come up with five features describing it (e.g., shown a cat, a participant may respond “is alive”). This dataset is publicly available (https://mariamh.shinyapps.io/dinolabobjects/), and its collection procedures along with name agreement data are described in full by Hovhannisyan et al.^31^. Each presented picture has a word label, which the present analyses using language models would leverage. For consistency, plural countable words were converted into singular forms; except for one case where the dataset included both a plural “scales” (referring to an animal’s skin) and “scale” (referring to a weight-measuring tool).

#### 4.1.2. Concept analysis

Our analyses focused on the fifty features most associated with the image concepts in a previously collected dataset of concept features (Hovhannisyan et al., 2021). As our analyses involved binary classification, we defined each object as having (1) a given feature if at least three people described the object as such (otherwise 0). The present analysis involved predicting whether each object concept has or does not have a given semantic feature using a linear SVM (C = 1 default) based on embeddings generated with Llama 3.2-3B (see embedding procedure below). For each feature’s classification, the data was balanced via random pruning, such that the numbers of examples with (1) or without (0) a given feature; no feature was associated with a majority of concepts, so pruning entailed omitting examples without a given feature. Balancing was expected to make the accuracy estimates more intuitive and allow clearer comparisons between features. Stratified 2-fold cross-validation was performed, and accuracy was averaged across 10 repetitions with different random pruning.

The LLM analyses involved submitting a short text for each concept into Llama 3.2-3B and extracting the residual stream activity. The short text was defined as “*A/An {object}*” or “*Some {objects}*” for uncountable singular words or for the one plural word in the dataset. LLMs are designed to process structured text, and incorporating a determiner (“a/an/some”) consistently increases performance. Specifically, adding this small grammatical construct increased classification accuracy for 45 out of the 50 properties tested using Llama 3.2-3B compared to using the capitalized word alone (“*{Object}*”); mean accuracy is 79.5% (with determiner) versus 77.4% accuracy (without determiner). This improvement constitutes roughly half of the LLM’s advantage relative to the word2vec accuracy of 75.6%.

After submitting the short texts to Llama 3.2-3B, the model’s residual stream activity was extracted for either singular layers or while concatenating the values across layers [4, 16), given 0-indexed counting. These residual stream activations constituted a concept embedding with 36,864 dimensions. For Experiment 1, the concept embedding was reduced to 300 dimensions using principal component analysis to ensure a fair comparison with word2vec, which uses 300 dimensions; without dimensionality reduction, the average feature accuracy for the layer 4-16 analysis is slightly higher (80.3%) compared to with reduction (79.5%). Analysis of each alternative transformer model (BERT, Llama 3.3-70b-Instruct, etc.) followed the same procedures. For BERT, the analyses used the final two layers’ activations for the embeddings, as this was found to maximize performance. For the other LLMs, which contained more layers than Llama 3.2-3B (e.g., Llama 3.3-70b contains 80 layers whereas Llama 3.2-3B has 28 layers), embeddings were defined based on layer depths proportional to those of Llama 3.2-3B (e.g., rather than layer 4, layer 11 was used for an 80-layer model).

For the word2vec comparison, we used pre-trained Word2Vec embeddings trained on Google News (300 dimensions) ^7^. Some object words were not available in the embeddings database, and these were manually changed (e.g., “*aeroplane*” was changed to “*airplane*” and “*candycane*” to just “*candy*”). Additionally, 27% of concepts’ labels spanned multiple words (e.g., “alarm clock”), and for these, the word2vec embeddings were computed separately for each word and then averaged. Note, the results do not meaningfully change if the analysis exclusively examines single-word concepts (mean Llama 3.2-3B accuracy = 78.9%, mean word2vec accuracy = 75.6%).

### 4.2. Four-stage experiment used for Studies 1B, 2A, and 2B

Study 1B (fMRI), Study 2A (behavioral), and Study 2B (fMRI) all use data collected from one four-stage experiment. For this experiment, 76 fluent English-speaking participants were recruited from the local community and screened for no history of neurological damage or mild cognitive impairment. Sixteen participants did not complete at least one of the four stages (e.g., withdrew after Stage 1), and these participants were not analyzed here. The final set of 60 participants included a younger adult subset (N = 35, M_age_ = 22.8 [SD = 3.3], 64% female, 36% male) and older adult subset (N = 31, M_age_ = 71.5 [SD = 4.5], 64% female, 36% male), although none of the present analyses focus on age-related differences. The research was approved by the Duke University Institutional Review Board.

This experiment consisted of four stages. Stages 1, 3, and 4 involve the presentation of object pictures or words in isolation, and these stages fMRI data were used for the Study 1B analysis. Stage 2 involved the presentation of a scene picture and then an object picture.

Participants’ behavioral responses, rating how likely it would be to find the presented object in the previously presented scene, were used for Study 2A. The fMRI data collected during this stage were used for Study 2B. These four stages are each detailed below.

In Stage 1, participants completed trials where they were shown an object (e.g., a tractor) with a label (e.g., “tractor”), and participants used a 4-point scale to rate how accurately the label described the object (1 = “*does not describe the object*”; 4 = “*exact description*”). The task was designed to elicit high ratings (mean rating = 3.60). One week later, participants completed the Stage 2 encoding task (see above) and were asked to return approximately 24hrs (range: 20-28 hours) to complete the Stage 3 and 4, conceptual and perceptual retrieval task, respectively.

For Stage 2, participants were shown a picture of a familiar scene (e.g., a farm) followed by a picture of an object (e.g., a tractor). Participants used a 4-point scale to report how likely it would be to find the object in the scene (1 = “*Very unlikely*”, 4 = “*Very likely*”). Each participant completed 114 trials. One-third of trials showed scene-object pairs designed to be incongruent (mean rating = 1.28), one-third showed pairs that were neither congruent nor incongruent (mean rating = 2.11), and one-third showed pairs designed to be congruent (mean rating = 3.67). The scene-object pairs were counterbalanced across participants, so each scene and object appeared in all three conditions, and there were 342 possible scene-object pairs in total.

In the Stage 3 conceptual retrieval task, participants were presented 144 words; 114 were the object labels corresponding to objects shown in the Stage 2 encoding task, and 30 were object labels representing new concepts. For each word, participants indicated whether they had previously seen the object concept during encoding using a 4-point scale (1 = “*definitely new*”, 2 = “*probably new*”, 3 = “*probably old*”, 4 = “*definitely old*”). Participants successfully recognized most objects, and the mean hit rate (3 or 4 response to old images) was 76%. The fMRI analyses would focus only on the 114 old trials, irrespective of participants’ responses.

In the Stage 4 visual retrieval task, participants were shown 126 images of objects. Among these, 96 images were of the exact objects shown in the Stage 2 encoding task, 18 images were perceptually similar lures of the original objects (e.g., a blue tractor Stage 2 depicted a red one), and 12 images represented new concepts not seen at the Stage 2 encoding task. Participants responded (i) “old”, (ii) “similar”, or (iii) “new” to each image, and successfully responded to most images, with a mean old hit rate of 64% and a mean similar hit rate of 49%. The fMRI analyses would focus on the 114 old or similar trials, again irrespective of participants’ responses. Preliminary tests showed that the inclusion of miss trials for the conceptual and visual RSA tasks did not influence the patterns of significance in the results.

### 4.3. fMRI collection and preprocessing

#### 4.3.1. MRI acquisition

During each of the four stages, MRI data were collected using a General Electric 3T MR750 scanner and an 8-channel head coil. Anatomical images were acquired using a T1-weighted echo-planar sequence (96 slices at 0.9×0.9×1.9 mm^3^). Functional images were acquired using an echo-planar imaging sequence (repetition time = 2000 ms, echo time = 30 ms, field of view = 19.2 cm, 36 oblique slices with voxel dimensions of 3×3×3 mm). Stimuli were projected onto a mirror at the back of the scanner bore, and responses were recorded using a four-button fiber-optic response box (Current Designs, Philadelphia, PA, USA). Functional resting-state images were collected from the participants using the same parameters (210 volumes, 7 minutes). The BOLD timeseries were resampled into standard space with a spatial resolution of 2×2×2 mm^3^ or 97×115×97 voxels.

The below descriptions of anatomical and functional preprocessing were automatically generated by fMRIPrep with the express intention that users should copy and paste this text into their manuscripts unchanged.

#### 4.3.2. Anatomical data preprocessing

A total of two T1-weighted (T1w) images were found within the input BIDS dataset. All of them were corrected for intensity non-uniformity (INU) with N4BiasFieldCorrection ^53^ distributed with ANTs 2.3.3.^54^ The T1w-reference was then skull-stripped with a Nipype implementation of the antsBrainExtraction.sh workflow (from ANTs), using OASIS30ANTs as a target template.

Brain tissue segmentation of cerebrospinal fluid (CSF), white-matter (WM), and gray-matter (GM) was performed on the brain-extracted T1w using fast (FSL 6.0.5.1).^55^ An anatomical T1w-reference map was computed after registration of two T1w images (after INU-correction) using mri_robust_template (FreeSurfer 7.3.2).^56^

Brain surfaces were reconstructed using recon-all (FreeSurfer 7.3.2),^57^ and the brain mask estimated previously was refined with a custom variation of the method to reconcile ANTs-derived and FreeSurfer-derived segmentations of the cortical gray-matter of Mindboggle.^58^ Volume-based spatial normalization to one standard space (MNI152NLin2009cAsym) was performed through nonlinear registration with antsRegistration (ANTs 2.3.3), using brain-extracted versions of both T1w reference and the T1w template. The following template was selected for spatial normalization and accessed with TemplateFlow (23.0.0):^59^ ICBM 152 Nonlinear Asymmetrical template version 2009c.^60^

#### 4.3.3. Functional data preprocessing

For each of the seven BOLD runs found per participant (across all tasks and sessions), the following preprocessing was performed. First, a reference volume and its skull-stripped version were generated using a custom methodology of fMRIPrep. Head-motion parameters with respect to the BOLD reference (transformation matrices, and six corresponding rotation and translation parameters) are estimated before any spatiotemporal filtering using MCFLIRT (FSL 6.0.5.1, Jenkinson et al. 2002). BOLD runs were slice-time corrected to 0.972s (0.5 of slice acquisition range 0.00s-1.94s) using 3dTshift from AFNI.^61^

The BOLD time series (including slice-timing correction when applied) were resampled onto their original, native space by applying the transforms to correct for head motion. These resampled BOLD time-series will be referred to as preprocessed BOLD in original space, or just preprocessed BOLD. The BOLD reference was then co-registered to the T1w reference using bbregister (FreeSurfer) which implements boundary-based registration.^62^ Co-registration was configured with six degrees of freedom.

Several confounding time series were calculated based on the preprocessed BOLD: framewise displacement (FD), DVARS, and three region-wise global signals. FD was computed using two formulations following Power et al. (absolute sum of relative motions)^63^ and Jenkinson (relative root mean squared displacement between affines).^64^ FD and DVARS are calculated for each functional run, both using their implementations in Nipype (following the definitions by Power et al.^63^). The three global signals are extracted within the CSF, the WM, and the whole-brain masks. Principal components are estimated after high-pass filtering the preprocessed BOLD time series (using a discrete cosine filter with 128s cut-off) for the two CompCor variants: temporal (tCompCor) and anatomical (aCompCor). tCompCor components are then calculated from the top 2% variable voxels within the brain mask. For aCompCor, three probabilistic masks (CSF, WM, and combined CSF+WM) are generated in anatomical space. The implementation differs from that of Behzadi et al. in that instead of eroding the masks by two pixels on BOLD space, a mask of pixels that likely contain a volume fraction of GM is subtracted from the aCompCor masks. This mask is obtained by dilating a GM mask extracted from the FreeSurfer’s aseg segmentation, and it ensures components are not extracted from voxels containing a minimal fraction of GM. Finally, these masks are resampled into BOLD space and binarized by thresholding at 0.99 (as in the original implementation). Components are also calculated separately within the WM and CSF masks. For each CompCor decomposition, the k components with the largest singular values are retained, such that the retained components’ time series are sufficient to explain 50% of variance across the nuisance mask (CSF, WM, combined, or temporal). The remaining components are dropped from consideration.

The head-motion estimates calculated in the correction step were also placed within the corresponding confounds file. The confound time series derived from head motion estimates and global signals were expanded with the inclusion of temporal derivatives and quadratic terms for each.^65^ Additional nuisance time series are calculated by means of principal components analysis of the signal found within a thin band (crown) of voxels around the edge of the brain, as proposed by Patriat, Reynolds, and Birn.^66^ The BOLD time series were resampled into standard space, generating a preprocessed BOLD run in MNI152NLin2009cAsym space. First, a reference volume and its skull-stripped version were generated using a custom methodology of fMRIPrep. All resampling can be performed with a single interpolation step by composing all the pertinent transformations (i.e., head-motion transform matrices, susceptibility distortion correction when available, and co-registrations to anatomical and output spaces). Gridded (volumetric) resampling was performed using antsApplyTransforms (ANTs), configured with Lanczos interpolation to minimize the smoothing effects of other kernels.^67^ Non-gridded (surface) resampling was performed using mri_vol2surf (FreeSurfer). Many internal operations of fMRIPrep use NiLearn 0.9.1,^68^ mostly within the functional processing workflow. For more details of the pipeline, see the section corresponding to workflows in fMRIPrep’s documentation.

#### 4.3.4. Single-trial activity modeling

Analyses measured each voxel’s BOLD response in each trial. This was performed with first-level general linear models using the Least Squares Separate approach by Mumford et al.,^69^, which involves fitting a separate regression for each trial. The regression included a boxcar signal spanning the object presentation period. The regressor’s resulting coefficient represents the trial’s BOLD response. The regression additionally included a boxcar signal covering the presentation time of all other stimuli (i.e., every other object and, for the Stage 2 analysis, every scene). The linear models also included six translation/rotation regressors, three other covariates for head motion (FD, DVARS, and RSMD), and covariates for mean global, white-matter, and cerebrospinal signals. For all four tasks of each of the 60 participants (240 scans), 114 three-dimensional beta coefficient volumes were defined. The voxelwise betas were then organized into 246 ROIs based on the Brainnetome Atlas. The ROIs cover all neocortical areas along with the hippocampus, amygdala, thalamus, and basal ganglia.

### 4.4. Study 1B concept representational similarity analysis

Concept representation in the brain was examined using RSA. This involved generating embeddings by submitting short “*An {object}*” texts to Llama 3.2-3B and concatenating the residual stream responses across layers [4, 16). This parallels the Study 1A concept analysis. For each participant’s embeddings, each dimension was z-scored across their 114 different concept embeddings. This was not necessary for the Study 1A analyses, as a given dimension’s mean across embeddings will not influence the support vector machines or regressions. However, for RSA in Study 1B, which involves correlations between trials’ embeddings, this will have an impact, and preliminary analyses demonstrated that this normalization bolsters RSA effects (for any model tested, e.g., Llama 3.2-3B RSA or the word2vec RSA).

For each participant and each of the three isolated-item task stages, an RSM was prepared by taking correlations between trials’ concept embeddings. A corresponding NSM was produced for each ROI via across-trial correlations between ROIs’ voxelwise activations. The first-level analysis of each participant’s data in each stage involved a second-order Spearman correlation between the associated RSM and each ROI’s NSM. For each participant, their second-order correlations were averaged across the three stages. Then, second-level analysis submitted each participant’s mean correlation to a one-sample t-test, evaluating whether the group-wide RSA effect significantly surpasses zero. For the analysis of the second-stage fMRI data, an analogous procedure was performed for scene concept-level RSA (“*A {scene}*” embeddings) along with scene-and-object RSA (averaging trials’ “*An {object}*” and “*A {scene}*” embeddings).

For **Figure 2C**, RSA was also performed for embeddings based on the residual streams of singular layers. Here, RSA was conducted for four large-scale areas, defined by the regional labels of the Brainnetome Atlas: the Occipital cortex (averaging the NSMs across medioventral and lateral occipital ROIs), the inferior temporal lobe (inferior temporal, fusiform, and parahippocampal ROIs), the parietal lobe (precuneus, inferior parietal lobule, superior parietal lobule ROIs), and the prefrontal cortex (orbitofrontal, inferior frontal, middle frontal, and superior frontal ROIs).

### 4.5. Study 2A relational behavioral analysis

The Study 2A analyses predicted the mean rating assigned to the 342 scene-object pairs using a ridge regression (α = 1 default), leave-one-out cross-validation, and clipping predictions to be between 1 and 4. To produce the LLM embeddings, two short texts corresponding to each pair were submitted to Llama 3.2-3B, and the residual stream activity was extracted for layers [4, 16) and concatenated to form an embedding with 36,864 dimensions. The two short texts consisted of “*At the {scene}, an {object}*” and “*An {object} at {the scene}*” with changes to the object determiner (“*a/an/some*”) or the scene proposition (“*at/in/*on”) as appropriate.

The determiners and propositions were prepared manually. Each object had a consistent determiner across all three of its scene-object pairs, and the same is true for each scene’s proposition. For the scene proposition, “at” was used as the default (96/114 scenes) and only changed for scenes where “at” seemed unnatural (e.g., “*On the balcony*” rather than “*At the balcony*”). Rather than just one text, two texts were used to capture both a scene → object (“*On the balcony, a violin*”) order and an object → scene order (“*A violin on the balcony*”). We presumed that this averaging would minimize potentially arbitrary order effects. The extracted residual stream values were averaged across the two texts to produce the relational embedding. Testing just texts’ embeddings individually shows that the “*At the {scene}, an {object}*” embeddings yield higher accuracy (R^2^ =.70) than the “*An {object} at {the scene}*” embeddings (R^2^ =.53); the averaged embedding accuracy is R^2^ =.67; see classification procedure below. Nonetheless, the average embedding across both scene/object orders was used as we expected this would increase the generalizability of the results. As with the concept classification, the text structure employed improves embedding quality. By comparison, averaging “*An {object} and {scene}*” and *“A {scene} and {object}*” produces R^2^ =.65, which is slightly lower. However, deviating from grammar entirely with “*{Object} {scene}*” and “*{Scene} {object}*” yields just R^2^ =.55, which is a starker decrease. For future research employing this methodology, these results suggest that, at minimum, the short texts should be grammatically proper.

For the word2vec comparisons, we measured the Pearson correlation between the object and scene word2vec embeddings. This technique has been used in prior work seeking an objective measure of two words’ relatedness.^70^ This scalar was then used as the sole predictor in a ridge regression with [1, 4] clipping and leave-one-out cross-validation, as in the LLM analysis. The regression predicted scene-object likelihood. This notably yields similar results (R^2^ =.365) as a Pearson correlation between similarity and scene-object likelihood (R^2^ =.368; Spearman ρ^2^ =.340). We explored predictions based on the difference between the scene and object word2vec embeddings, producing a 300-dimension difference vector. However, this difference vector did not yield above-chance predictions (R^2^ < 0), nor did an absolute difference vector (R^2^ < 0).

We additionally report brief analyses using a toy dataset involving 20 animals (10 herbivores and 10 carnivores) and 20 foods (10 plants and 10 meats) used to prepare texts “*An {animal} and a {food}*” and labeled based on whether the animal would eat the food. Classification using similar procedures as above yielded accuracy well above chance (∼70%) with group-2-fold cross-validation, training on herbivore texts and testing on carnivore texts or vice versa. These results are, however, just meant as an illustration to provide intuition, and prior studies provide more formal decompositions of LLMs’ residual streams into features^25^

### 4.6. Study 2B relation-level representational similarity analysis

#### 4.6.1. Measuring relational representation

Relational representation was similarly assessed using RSA, now using relational embeddings. These embeddings were generated by averaging the residual stream layer [4, 16) responses across the texts “*At the {scene}, an {object}*” and “*An {object} at the {scene}*”, which parallels the Study 2A relational analysis. Across the study, each object was associated with three pairs of text – e.g., for an apple, there is “*At the orchard, an apple*”, “*At the gym, an apple*”, “*At the iceberg, an apple*”. To remove concept-level information about the apple from each of these embeddings, the three {scene}-apple embeddings were averaged, and this average was subtracted from the other three embeddings. For example, by subtracting this mean from the “At the orchard, an apple” embedding, the resulting object-normalized embedding now just expresses the orchard-apple relation and the orchard concept. To remove the orchard concept information, the object-normalized embedding mean was computed for “*At the orchard, an apple*”, “*At the orchard, a Ferris wheel*”, and “*At the orchard, a police car*”. This mean was then subtracted from the object-normalized embedding of “*At the orchard, an apple*” producing an embedding that has omitted both object and scene concept-level information.

Using this relational embedding, RSA was performed. First, to confirm that this embedding omitted effectively all concept-level information, RSA was attempted while applying these relational embeddings to analyze the fMRI data of the isolated-item task stage – i.e., stages where no scenes were shown and thus RSA should yield null results. Afterward, analyses turned to the second stage’s fMRI data. Here, group-wide trends were evaluated using both one-sample t-tests relative to zero and paired t-test comparing the relational RSA effect to the object-and-scene concept RSA effect.

#### 4.6.2. Predicting normative fit

For the final analyses predicting the normative fit of participants’ responses in the scene-object task, single-trial measures of representational strength were computed using the procedures by Davis et al.^12^. Specifically, a row-by-row correlation was computed between each row of the (114 × 114) relational-embedding RSM and the (114 × 114) NSM, producing 114 values describing representational strength in each trial. For each participant, their representational strength values were correlated with their normative fit scores – that is, the absolute difference between the response and 4, 2.5, or 1, depending on whether a given trial was in the incongruent, neutral, or congruent condition (see task design above). For the group-level analysis, the Pearson correlations were submitted to a one-sample t-test; note that this is largely equivalent to a multilevel regression of “*accuracy ∼ 1 + representation + (1 + representation | participant)*”.

## Data availability

The data collected will be shared upon request via email to the corresponding author, Paul C. Bogdan, or principal investigators Roberto Cabeza and Simon W. Davis. We have been explicitly told by our IRB that we do not have permission to upload the collected data to a public repository.

## References

1. Nelson, D. L., McEvoy, C. L. & Schreiber, T. A. The University of South Florida free association, rhyme, and word fragment norms. Behavior Research Methods, Instruments, & Computers 36, 402–407 (2004).

2. Hebart, M. N., Zheng, C. Y., Pereira, F. & Baker, C. I. Revealing the multidimensional mental representations of natural objects underlying human similarity judgements. Nature human behaviour 4, 1173–1185 (2020).

3. Taylor, K. I., Moss, H. E. & Tyler, L. K. The conceptual structure account: A cognitive model of semantic memory and its neural instantiation. Neural basis of semantic memory 265–301 (2007).

4. Shen, X. et al. Using connectome-based predictive modeling to predict individual behavior from brain connectivity. nature protocols 12, 506–518 (2017).

5. Pereira, F., Mitchell, T. & Botvinick, M. Machine learning classifiers and fMRI: a tutorial overview. Neuroimage 45, S199–S209 (2009).

6. Haxby, J. V. Multivariate pattern analysis of fMRI: the early beginnings. Neuroimage 62, 852– 855 (2012).

7. Mikolov, T., Chen, K., Corrado, G. & Dean, J. Efficient estimation of word representations in vector space. arXiv preprint arXiv:1301.3781 (2013).

8. Kriegeskorte, N., Mur, M. & Bandettini, P. A. Representational similarity analysis-connecting the branches of systems neuroscience. Frontiers in systems neuroscience 2, 4 (2008).

9. Waltz, J. A. et al. A system for relational reasoning in human prefrontal cortex. Psychological science 10, 119–125 (1999).

10. Waltz, J. A. et al. Relational integration and executive function in Alzheimer’s disease. Neuropsychology 18, 296 (2004).

11. Clarke, A. & Tyler, L. K. Object-specific semantic coding in human perirhinal cortex. Journal of Neuroscience 34, 4766–4775 (2014).

12. Davis, S. W. et al. Visual and semantic representations predict subsequent memory in perceptual and conceptual memory tests. Cerebral Cortex 31, 974–992 (2021).

13. Martin, C. B., Douglas, D., Newsome, R. N., Man, L. L. & Barense, M. D. Integrative and distinctive coding of visual and conceptual object features in the ventral visual stream. elife 7, e31873 (2018).

14. Mayes, A., Montaldi, D. & Migo, E. Associative memory and the medial temporal lobes. Trends in cognitive sciences 11, 126–135 (2007).

15. Hannula, D. E. & Ranganath, C. Medial temporal lobe activity predicts successful relational memory binding. Journal of Neuroscience 28, 116–124 (2008).

16. Eichenbaum, H., Yonelinas, A. P. & Ranganath, C. The medial temporal lobe and recognition memory. Annu. Rev. Neurosci. 30, 123–152 (2007).

17. Hobeika, L., Diard-Detoeuf, C., Garcin, B., Levy, R. & Volle, E. General and specialized brain correlates for analogical reasoning: A meta-analysis of functional imaging studies. Human brain mapping 37, 1953–1969 (2016).

18. Fresnoza, S. & Ischebeck, A. Probing our built-in calculator: A systematic narrative review of noninvasive brain stimulation studies on arithmetic operation-related brain areas. Eneuro 11, (2024).

19. Ackerman, C. M. & Courtney, S. M. Spatial relations and spatial locations are dissociated within prefrontal and parietal cortex. Journal of Neurophysiology 108, 2419–2429 (2012).

20. Beiser, D. G. & Houk, J. C. Model of cortical-basal ganglionic processing: encoding the serial order of sensory events. Journal of Neurophysiology 79, 3168–3188 (1998).

21. Wertheim, J. & Ragni, M. The neural correlates of relational reasoning: a meta-analysis of 47 functional magnetic resonance studies. Journal of cognitive neuroscience 30, 1734–1748 (2018).

22. Hélie, S., Ell, S. W. & Ashby, F. G. Learning robust cortico-cortical associations with the basal ganglia: an integrative review. Cortex 64, 123–135 (2015).

23. Morin, T. M., Moore, K. N., Isenburg, K., Ma, W. & Stern, C. E. Functional reconfiguration of task-active frontoparietal control network facilitates abstract reasoning. Cerebral Cortex 33, 5761–5773 (2023).

24. Watson, C. E. & Chatterjee, A. A bilateral frontoparietal network underlies visuospatial analogical reasoning. Neuroimage 59, 2831–2838 (2012).

25. Bricken, T., et al. Towards monosemanticity: Decomposing language models with dictionary learning. Transformer Circuits Thread 2, (2023).

26. Kumar, S. et al. Shared functional specialization in transformer-based language models and the human brain. Nature communications 15, 5523 (2024).

27. Franch, M. et al. A vectorial code for semantics in human hippocampus. bioRxiv 2025–02 (2025).

28. Mischler, G., Li, Y. A., Bickel, S., Mehta, A. D. & Mesgarani, N. Contextual feature extraction hierarchies converge in large language models and the brain. Nature Machine Intelligence 1– 11 (2024).

29. Zhang, X., Wang, S., Lin, N., Zhang, J. & Zong, C. Probing word syntactic representations in the brain by a feature elimination method. in vol. 36 11721–11729 (2022).

30. Ravfogel, S., Elazar, Y., Gonen, H., Twiton, M. & Goldberg, Y. Null it out: Guarding protected attributes by iterative nullspace projection. arXiv preprint arXiv:2004.07667 (2020).

31. Hovhannisyan, M. et al. The visual and semantic features that predict object memory: Concept property norms for 1,000 object images. Memory & Cognition 49, 712–731 (2021).

32. Dixon, M. L. et al. Heterogeneity within the frontoparietal control network and its relationship to the default and dorsal attention networks. Proceedings of the National Academy of Sciences 115, E1598–E1607 (2018).

33. Bogdan, P. C., Iordan, A., Shobrook, J. & Dolcos, F. ConnSearch: A framework for functional connectivity analyses designed for interpretability and effectiveness at limited sample sizes. NeuroImage 278, 120274 (2023).

34. Noonan, K. A., Jefferies, E., Visser, M. & Lambon Ralph, M. A. Going beyond inferior prefrontal involvement in semantic control: evidence for the additional contribution of dorsal angular gyrus and posterior middle temporal cortex. Journal of cognitive neuroscience 25, 1824–1850 (2013).

35. Hadj-Bouziane, F., Meunier, M. & Boussaoud, D. Conditional visuo-motor learning in primates: a key role for the basal ganglia. Journal of Physiology-Paris 97, 567–579 (2003).

36. Wymbs, N. F., Bassett, D. S., Mucha, P. J., Porter, M. A. & Grafton, S. T. Differential recruitment of the sensorimotor putamen and frontoparietal cortex during motor chunking in humans. Neuron 74, 936–946 (2012).

37. Howard, C. M., Huang, S., Hovhannisyan, M., Cabeza, R. & Davis, S. W. Differential Mnemonic Contributions of Cortical Representations during Encoding and Retrieval. Journal of Cognitive Neuroscience 36, 2137–2165 (2024).

38. Suzuki, W. A. & Naya, Y. The perirhinal cortex. Annual review of neuroscience 37, 39–53 (2014).

39. Liuzzi, A. G. et al. Left perirhinal cortex codes for semantic similarity between written words defined from cued word association. Neuroimage 191, 127–139 (2019).

40. Price, A. R., Bonner, M. F., Peelle, J. E. & Grossman, M. Converging evidence for the neuroanatomic basis of combinatorial semantics in the angular gyrus. Journal of Neuroscience 35, 3276–3284 (2015).

41. Vann, S. D., Aggleton, J. P. & Maguire, E. A. What does the retrosplenial cortex do? Nature reviews neuroscience 10, 792–802 (2009).

42. Epstein, R. A. Parahippocampal and retrosplenial contributions to human spatial navigation. Trends in cognitive sciences 12, 388–396 (2008).

43. Thornton, M. A. & Tamir, D. I. Neural representations of situations and mental states are composed of sums of representations of the actions they afford. Nature communications 15, 620 (2024).

44. Wutz, A., Loonis, R., Roy, J. E., Donoghue, J. A. & Miller, E. K. Different levels of category abstraction by different dynamics in different prefrontal areas. Neuron 97, 716–726 (2018).

45. Jefferies, E. & Lambon Ralph, M. A. Semantic impairment in stroke aphasia versus semantic dementia: a case-series comparison. Brain 129, 2132–2147 (2006).

46. Noonan, K. A., Jefferies, E., Visser, M. & Lambon Ralph, M. A. Going beyond inferior prefrontal involvement in semantic control: evidence for the additional contribution of dorsal angular gyrus and posterior middle temporal cortex. Journal of cognitive neuroscience 25, 1824–1850 (2013).

47. Whitney, C., Kirk, M., O’Sullivan, J., Lambon Ralph, M. A. & Jefferies, E. The neural organization of semantic control: TMS evidence for a distributed network in left inferior frontal and posterior middle temporal gyrus. Cerebral cortex 21, 1066–1075 (2011).

48. Snowden, J. S. et al. Semantic dementia and the left and right temporal lobes. Cortex 107, 188–203 (2018).

49. Gainotti, G. & Marra, C. Differential contribution of right and left temporo-occipital and anterior temporal lesions to face recognition disorders. Frontiers in Human Neuroscience 5, 55 (2011).

50. Mink, J. W. The basal ganglia: focused selection and inhibition of competing motor programs. Progress in neurobiology 50, 381–425 (1996).

51. Humphries, M. D. & Prescott, T. J. The ventral basal ganglia, a selection mechanism at the crossroads of space, strategy, and reward. Progress in neurobiology 90, 385–417 (2010).

52. Nir-Cohen, G., Kessler, Y. & Egner, T. Neural substrates of working memory updating. Journal of Cognitive Neuroscience 32, 2285–2302 (2020).

53. Tunison, E., Sylvain, R., Sterr, J., Hiley, V. & Carlson, J. M. No money, no problem: enhanced reward positivity in the absence of monetary reward. Frontiers in human neuroscience 13, 41 (2019).

54. Avants, B. B., Epstein, C. L., Grossman, M. & Gee, J. C. Symmetric diffeomorphic image registration with cross-correlation: evaluating automated labeling of elderly and neurodegenerative brain. Medical image analysis 12, 26–41 (2008).

55. Zhang, Y., Brady, M. & Smith, S. Segmentation of brain MR images through a hidden Markov random field model and the expectation-maximization algorithm. IEEE transactions on medical imaging 20, 45–57 (2001).

56. Reuter, M., Rosas, H. D. & Fischl, B. Highly accurate inverse consistent registration: a robust approach. Neuroimage 53, 1181–1196 (2010).

57. Dale, A. M., Fischl, B. & Sereno, M. I. Cortical surface-based analysis: I. Segmentation and surface reconstruction. Neuroimage 9, 179–194 (1999).

58. Klein, A. et al. Mindboggling morphometry of human brains. PLoS computational biology 13, e1005350 (2017).

59. Ciric, R. et al. TemplateFlow: FAIR-sharing of multi-scale, multi-species brain models. Nature Methods 19, 1568–1571 (2022).

60. Fonov, V. S., Evans, A. C., McKinstry, R. C., Almli, C. R. & Collins, D. Unbiased nonlinear average age-appropriate brain templates from birth to adulthood. NeuroImage 47, S102 (2009).

61. Cox, R. W. & Hyde, J. S. Software tools for analysis and visualization of fMRI data. NMR in Biomedicine: An International Journal Devoted to the Development and Application of Magnetic Resonance In Vivo 10, 171–178 (1997).

62. Greve, D. N. & Fischl, B. Accurate and robust brain image alignment using boundary-based registration. Neuroimage 48, 63–72 (2009).

63. Power, J. D. et al. Methods to detect, characterize, and remove motion artifact in resting state fMRI. Neuroimage 84, 320–341 (2014).

64. Jenkinson, M., Bannister, P., Brady, M. & Smith, S. Improved optimization for the robust and accurate linear registration and motion correction of brain images. Neuroimage 17, 825–841 (2002).

65. Satterthwaite, T. D. et al. An improved framework for confound regression and filtering for control of motion artifact in the preprocessing of resting-state functional connectivity data. Neuroimage 64, 240–256 (2013).

66. Patriat, R., Reynolds, R. C. & Birn, R. M. An improved model of motion-related signal changes in fMRI. Neuroimage 144, 74–82 (2017).

67. Lanczos, C. Evaluation of noisy data. Journal of the Society for Industrial and Applied Mathematics, Series B: Numerical Analysis 1, 76–85 (1964).

68. Abraham, A. et al. Machine learning for neuroimaging with scikit-learn. Frontiers in neuroinformatics 8, 14 (2014).

69. Mumford, J. A., Turner, B. O., Ashby, F. G. & Poldrack, R. A. Deconvolving BOLD activation in event-related designs for multivoxel pattern classification analyses. Neuroimage 59, 2636– 2643 (2012).

70. Walsh, C. R. & Rissman, J. Behavioral representational similarity analysis reveals how episodic learning is influenced by and reshapes semantic memory. Nature Communications 14, 7548 (2023).

